# A phage-encoded host transcription modulator, Htm, reprograms the transcriptome of ESBL *Escherichia coli*

**DOI:** 10.64898/2026.10.06.757011

**Authors:** Saar S. F. Van Overfelt, Viktor Hundtofte Mebus, Jonathan Brewer, Rob Lavigne, Anders Nørgaard Sørensen, Lone Brøndsted

## Abstract

During phage infection, phages reprogram their host to secure energy, building blocks for their genome, and proteins, as well as the transcription and translation machinery required to replicate their genome and produce phage particles. This also includes reprogramming the host transcriptome to modify host gene expression and align it with the phage’s needs during infection. In this study, we present a putative early-expressed protein, Gp204 of *Mosigvirus* AV110, renamed Host transcription modulator (Htm), that has a broad impact on the extended-spectrum β-lactamase (ESBL)-associated *E. coli* transcriptome. Overexpression of Htm decreased the growth rate of ESBL-*E. coli* and caused DNA condensation. Htm is structurally similar to bacterial enhancer-binding proteins, binds ssDNA, and alters the expression levels of over 700 *E. coli* genes when expressed. The majority of these genes were upregulated and are involved in multiple pathways, such as galactose metabolism and pentose and glucuronate interconversions, which may promote the synthesis of purines, pyrimidines, and histidine, all of which are required for the replication, transcription, and translation of the phage genome. Furthermore, energy production is likely to be upregulated by increased availability of acetyl-CoA, resulting from, among other factors, fatty acid degradation. Htm also downregulated the expression of the structural genes involved in flagellar assembly and multiple two-component systems. In conclusion, Htm could promote phage progeny production by increasing the availability of nucleotides, amino acids, and energy, while inhibiting non-essential processes, such as flagella formation.

**Importance:** Early-expressed proteins have a central role during phage infection by rapidly reprogramming a wide range of host cellular processes. However, the function and impact on the host of many of these proteins remain poorly understood. Investigating these proteins provides fundamental insights into phage-host co-evolution and the mechanisms phages use to support their successful infection and progeny formation. In a broader context, this knowledge may contribute to the development of antimicrobials with a novel mode of action against multidrug-resistant bacteria, such as ESBL *E. coli* or biotechnological tools to precisely alter bacterial physiology. Our findings indicate that the phage-encoded protein Htm (Host transcription modulator) acts as a regulator of host transcription and demonstrate how early-expressed phage proteins can take control of the host.

## Introduction

Bacteriophages (phages) are bacterial viruses that depend entirely on their hosts to complete their life cycles. Unlike bacteria, phages have a relatively small genome that encodes only a limited number of essential functions, and they lack the capacity to perform most of the processes required to replicate their genome and produce proteins (1). Therefore, phages must exploit the host’s resources to propagate (2). This includes the use of the transcription and translation machinery to produce viral proteins, as well as the pool of nucleotides, amino acids, and energy required to replicate their genome and assemble new virions (2).

For their transcription, phages can either encode their own RNA polymerase or rely partly or entirely on the host’s RNA polymerase. Phage T7 requires the *E. coli* RNA polymerase to transcribe its early genes but encodes its own polymerase for the transcription of the middle and late genes (3). To redirect the host RNA polymerase, its early promoters contain strong recognition sites for the σ^70^-bound RNA polymerase, which is sufficient to drive transcription of those genes. Other phages, like T4, depend on the host’s RNA polymerase to transcribe all phage genes. Similar to T7 early promoters, T4 early promoters contain all elements for efficient binding by the σ^70^-bound RNA polymerase, but ADP-ribosylation of one α subunit of the RNA polymerase is necessary to further enhance transcription of the early genes (4, 5). Afterward, the host RNA polymerase is directed toward the middle promoters through binding with the early phage proteins MotA, which binds σ^70^ and the MotA-box in the middle promoters, and AsiA, which binds σ^70^, thereby preventing host transcription (6, 7). The host machinery is also needed for translation and is modified to support a more efficient infection. During phage T4 infection, ModB attaches an RNA chain to the ribosomal proteins L2 and S1, thereby increasing the burst size and accelerating lysis (8).

Phages also need resources from their host, primarily energy, nucleotides, and amino acids, to support their propagation. These resources can be obtained through various strategies. First, the host’s consumption of energy and nucleotides can be reduced by inhibiting host transcription. Phage T7 achieves this via the early-expressed protein Ocr, which binds σ^70^ of the RNA polymerase and inhibits its activity (9). Phage T4 encodes Alc, leading to transcription termination (10). Second, nucleotide levels in the cell can be increased by degrading the host mRNAs and genome early in infection (11–13). During T4 infection, this is achieved through the activity of Srd, which enhances host RNase E, and the endonucleases EndoII and EndoIV (14, 15). At the same time, this inhibits transcription and translation of host proteins. Finally, host metabolic processes can be altered to promote nucleotide and amino acid biosynthesis, as well as energy production or conservation. However, the effect of phage infection on the host metabolome varies greatly depending on the phage (16). For example, infection with phage T4 results in increased expression of genes involved in energy production and carbohydrate metabolism, while reducing ribosome biogenesis and amino acid metabolism (12). Conversely, infection of *P. aeruginosa* with phage JBD44 alters nitrogen metabolism, two-component systems, and biofilm formation (17). These metabolic changes can be achieved by changing gene expression levels in these pathways via phage-encoded transcriptional regulators (18, 19), although our understanding of how individual phage proteins influence the host transcriptome and metabolome remains limited.

In a previous screening of early-expressed proteins of *Mosigvirus* AV110, Gp204 was identified as an inhibitor of the growth of ESBL-producing *E. coli* (20). Here, we aimed to determine the function of Gp204 and its effect on specific cellular processes and the transcriptome of ESBL *E. coli* through microscopy and RNA sequencing. Our results show that Gp204, renamed Host transcription modulator (Htm), alters the host transcriptome and influences the expression of genes involved in a wide range of processes. As a result, Htm promotes nucleotide and nucleotide sugar metabolism, phosphotransferase systems, and processes involved in energy supply, but reduces cell motility and certain two-component systems.

## Results

### Gp204 inhibits the growth of *E. coli*

The protein Gp204 of *Mosigvirus* AV110 was previously identified as an early-expressed protein that inhibits the growth of ESBL098 (20). To verify whether the growth-inhibitory effect of Gp204 is limited to ESBL098, it was expressed in a genetically diverse panel of ESBL/AmpC *E. coli* strains and *E. coli* Stellar™. Gp204 effectively reduced the growth of all tested strains by approximately four to five logs, but smaller, potentially resistant colonies were seen in all dilutions (Figure 1A). When following the growth of ESBL098 pPS26::*gp204* in the presence and absence of rhamnose by OD600 measurements, it was observed that Gp204 did not completely inhibit growth but did strongly reduce the growth rate (Figure 1B). After thirteen hours, the growth rate increased again; however, in one of three replicates, this increase was observed after nine hours and was greater than in the other two. It is possible that a mutation occurred in this replicate, resulting in resistance to Gp204 and, therefore, faster growth. Using time-lapse microscopy, growth could be observed for approximately two hours after induction. Although growth stopped after two hours, the majority of the cells maintained their rod shape and appeared similar to the control for ten to twenty hours after induction (Video S1). There was some variation in the timing between the different replicates; however, in all replicates, after ten to twenty hours, the appearance of the cells gradually became more irregular, and the cells became smaller. In all replicates, the regrowth of resistant colonies was observed. Comparing colony sizes of ESBL098 transformed with the empty vector control and Gp204 showed that expressing Gp204 led to a slightly smaller colony area (Figure 1C). However, live-dead staining revealed no differences in the level of cell death (Area Under Curve: pPS26 = 32341 ± 4169, Gp204 = 33243 ± 4230, p = 0.81; Figure 1C). Overall, this indicates that Gp204 inhibits the growth of diverse ESBL/AmpC *E. coli* strains, likely targeting a conserved process without affecting cell shape or causing cell death.

**Figure 1:**
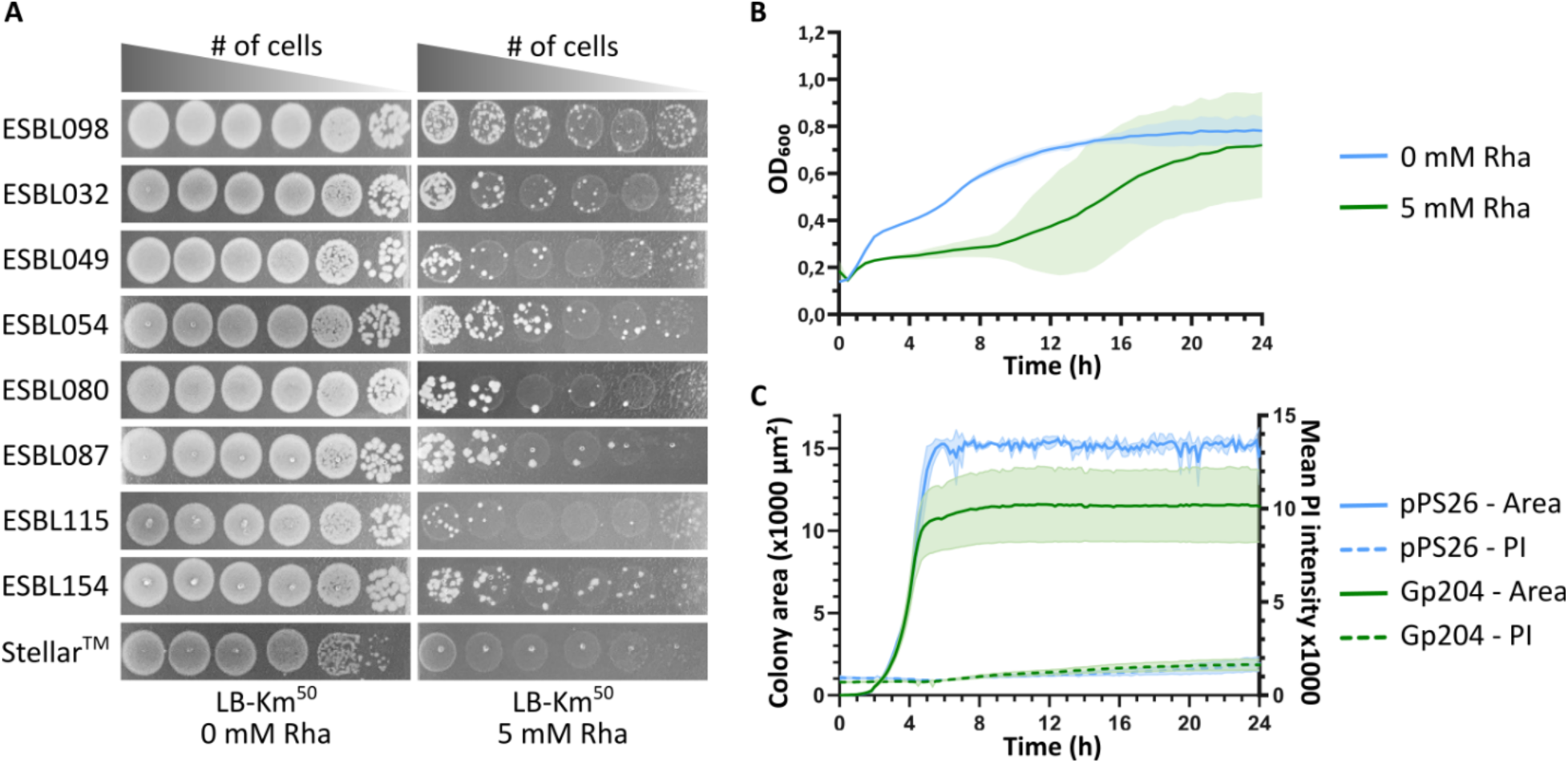
Gp204 inhibits the growth of *E. coli*. A) Tenfold dilutions of different ESBL/AmpC *E. coli* strains and *E. coli* Stellar^TM^ transformed with pPS26::*gp204* were spotted on LA plates without and with 5 mM rhamnose. B) The growth of ESBL098 pPS26::*gp204* was followed over time in the presence and absence of 5 mM rhamnose. The average and standard deviation of biological triplicates are shown. C) The measured colony area and mean propidium iodide (PI) intensity during time-lapse microscopy with live-dead staining of ESBL098 transformed with either the empty vector pPS26 or pPS26::*gp204* is shown. The average and standard deviation of biological triplicates are shown.

### Gp204 causes DNA condensation in *E. coli*

To better understand the cellular processes affected by Gp204, fluorescent microscopy was performed at one, two, and four hours of expressing Gp204, with staining the cell membrane, sites of active peptidoglycan synthesis, and DNA. After one hour, no differences were observed compared to the empty vector control (Figure S1A). After two and four hours, multiple cells showed compact nucleoid staining in the middle of the cell, suggesting DNA condensation. Additionally, septum formation was no longer initiated, although a limited number of cells had a fully formed septum (Figure 2A and Figure S1B). Peptidoglycan synthesis and the cell membrane remained unaffected by Gp204. Next, Gp204 localization in the cell was examined after four hours of expression through the binding of a fluorescent ligand to a HaloTag that was C-terminally fused to Gp204. This showed that Gp204 was mainly found at the cell poles, either on one or both poles (Figure 2B). A heatmap displaying Gp204 and DNA intensity distribution within the cell indicated that Gp204 is located at the cell poles. The Gp204 signal partially overlaps with the DNA at the cell poles, although it can also be located in the cytoplasm (Figure 2C). These findings demonstrate that Gp204 influences DNA without affecting cell morphology.

**Figure 2:**
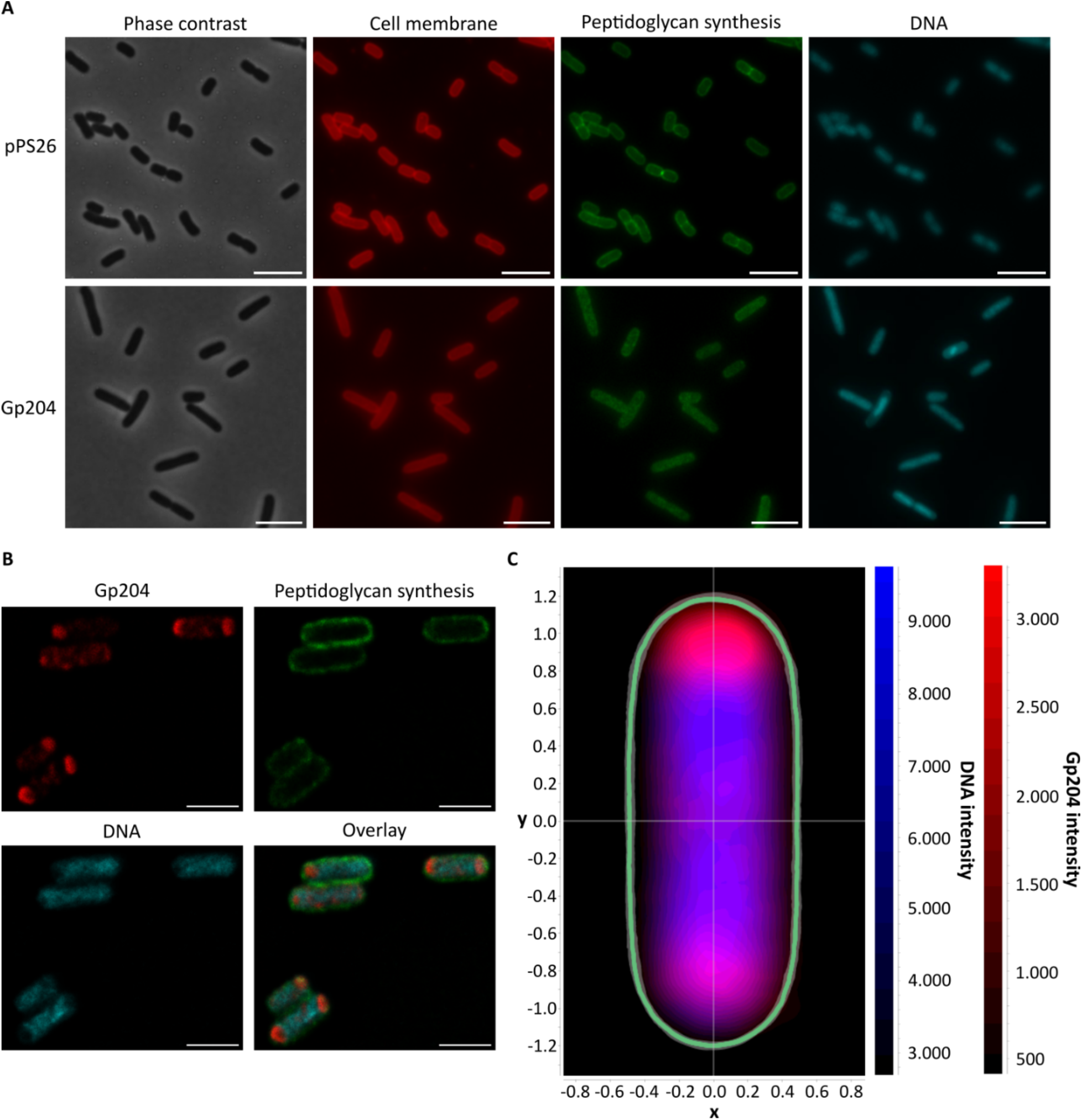
Gp204 is localized at the cell poles and causes DNA condensation. A) Phase contrast and fluorescence microscopy of ESBL098 transformed with pPS26 or pPS26::*gp204* after two hours of induction with staining of the cell membrane, peptidoglycan synthesis, and DNA. Representative images from fifteen images per condition are shown. B) Fluorescent images of ESBL098 pPS26::*gp204* after four hours of Gp204 expression showing the location of Gp204, peptidoglycan synthesis, DNA, and an overlay of all three. C) Heatmap of the DNA and Gp204 intensity distribution in the cell (n = 157 cells).

### Gp204 potentially functions as a transcriptional regulator

Next, we conducted *in silico* analyses to better understand the function of Gp204 through its protein structure and structural similarities to known proteins. The protein structure was predicted using AlphaFold 3 (21), resulting in a confident prediction (pTM = 0.75; Figure 3A). A conserved domain search identified a conserved AAA+ ATPase domain, characterized by a parallel β sheet and two α helices on one side and three on the other. This domain usually forms a hexamer (22). To test if Gp204 can also form a hexamer, its multimerization was predicted using the ipTM score from AlphaFold 3 predictions of different multimers, showing a peak with an ipTM score of 0.57 for the hexameric conformation (Figure 3B and Figure S2). However, this score remained just below 0.6, which is the lower threshold of the gray zone where no definitive conclusions can be made about the prediction’s accuracy. The ATPase domain of Gp204 contains the conserved Walker A motif, involved in ATP binding (22), but has a mutation in the Walker B motif. This motif is involved in ATP hydrolysis and consists of four hydrophobic residues followed by aspartic acid and glutamic acid, where aspartic acid is important for Mg^2+^ binding and glutamic acid for the catalytic activity (22). In Gp204, the glutamic acid is replaced by a second aspartic acid (22). Additionally, an insertion was found in the conserved ATPase domain, forming a small α-helix between α3 and β4.

**Figure 3:**
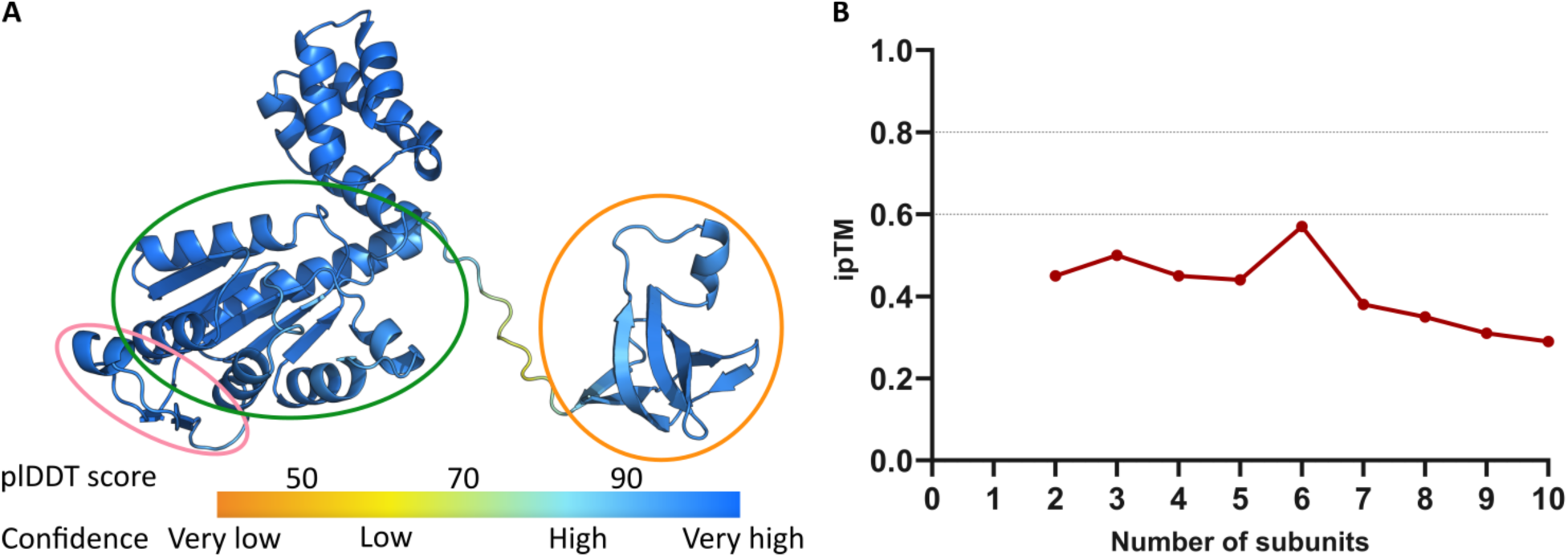
Gp204 encodes an AAA+ ATPase domain. A) The protein structure of Gp204 was predicted by AlphaFold 3 (21) and is colored according to the plDDT score of each amino acid.The AAA+ ATPase domain is shown in green, the insertion in the ATPase domain in pink and the N-terminal domain in orange. B) The multimerization of Gp204 was predicted based on the ipTM score of the corresponding AlphaFold 3 models, showing a peak for the hexameric conformation.

Comparing the monomeric structure of Gp204 to other protein structures with FoldSeek (23) showed that there is some structural similarity to transcriptional regulators, such as RtcR, PspF, and NtrC, but this similarity does not include the 80 amino acids of the N-terminal domain (Table **1** and Figure S3). These proteins, also known as bacterial enhancer binding proteins (bEBPs), regulate transcription by activating the σ^54^-bound RNA polymerase (24). The motif GAFTGA, located on a loop inserted in α3 of the conserved AAA+ ATPase domain structure, is important for σ^54^ interactions (25), however, Gp204 does not contain this loop or motif. The C-terminal domain of bEBPs interacts with enhancer binding sites through a helix-turn-helix motif (24), but is not present in Gp204 (Figure S3). Alternatively, Gp204 has an N-terminal domain that is not found in bEBPs but is present in other phage proteins. However, none of these proteins have been functionally annotated. A Foldseek search of the Gp204 N-terminal domain revealed that it has similarity to the C-terminal domain of NusG, which interacts with the transcription terminator ρ and the ribosomal protein S10 (26, 27).

**Table 1:** Overview of the FoldSeek results using the predicted monomeric structure of Gp204 in the 3Di/AA mode against the AFDB-PROTEOME and PBD database.

| AFDB/PDB ID | Description | Scientific name | Prob. | Seq. Id. | E-value |
| --- | --- | --- | --- | --- | --- |
| **AFDB-PROTEOME** |  |  |  |  |  |
| Q8ZN32 | Transcriptional regulator of two-component regulator protein, yfhA | *Salmonella enterica* subsp. enterica serovar Typhimurium str. LT2 | 1.00 | 14.8 | 1.17e-6 |
| Q32D18 | 2-component transcriptional regulator, yfhA | *Shigella dysenteriae* Sd197 | 1.00 | 15.2 | 4.01e-6 |
| P38035 | Transcriptional regulatory protein, rtcR | *Escherichia coli* K-12 | 1.00 | 10.7 | 2.99e-6 |
| Q8ZP59 | Transcription activator, pspF | *Salmonella enterica* subsp. enterica serovar Typhimurium str. LT2 | 1.00 | 13 | 6.81e-6 |
| P37344 | Psp operon transcriptional activator, pspF | *Escherichia coli* K-12 | 1.00 | 13.6 | 1.03e-5 |
| Q32F59 | Psp operon transcriptional activator, pspF | *Shigella dysenteriae* Sd197 | 1.00 | 11.9 | 8.60e-6 |
| A0A077ZED7 | DNA-binding transcriptional regulator, ntrC | *Trichuris trichiura* | 1.00 | 14.3 | 1.03e-5 |
| A0A0H3GGU1 | DNA-binding transcriptional regulator, ntrC | *Klebsiella pneumoniae* subsp. pneumoniae HS11286 | 1.00 | 14 | 1.64e-5 |
| Q9HVK3 | Transcriptional regulator, rtcR | *Pseudomonas aeruginosa* PAO1 | 1.00 | 12.2 | 7.65e-6 |
| A0A0H3GS68 | Phage shock protein operon transcriptional activator | *Klebsiella pneumoniae* subsp. pneumoniae HS11286 | 1.00 | 12 | 2.33e-5 |
| **PDB** |  |  |  |  |  |
| 5c3c | CbbQ/NirQ/NorQ domain protein | *Halothiobacillus neapolitanus* | 1.00 | 11.8 | 1.70e-5 |
| 2c96 | Psp operon transcriptional activator | *Escherichia coli* K-12 | 1.00 | 14.1 | 7.46e-6 |
| 1j7k | Holliday junction DNA helicase RuvB | *Thermotoga maritima* | 1.00 | 15.4 | 3.24e-5 |
| 2c98 | Psp operon transcriptional activator | *Escherichia coli* K-12 | 1.00 | 13.8 | 6.17e-5 |
| 1ny5 | Transcriptional regulator (NtrC family) | *Aquifex aeolicus* | 1.00 | 13.6 | 2.71e-5 |

### Gp204 has affinity for ssDNA

Due to the similarity of Gp204 to transcriptional regulators, its potential to bind ssDNA or dsDNA was tested in an electrophoretic mobility shift assay (EMSA) with the phage phiX174 genome in the single-stranded virion or double-stranded replicative form as the DNA templatee. The T4 ssDNA binding protein Gp32 was used as a positive control in combination with the phiX174 virion DNA, showing an upwards shift of the DNA band (Figure 4). Gp204 was purified from *E. coli* BL21 using a C-terminal His-tag, although this was complicated due to the inhibitory effect of Gp204 on *E. coli* BL21, resulting in a relatively low protein concentration. Loading the purified Gp204 without any DNA showed that DNA did not copurify with the protein or that the DNA concentration was below the detection limit. The EMSA showed that Gp204 does not bind to dsDNA but exhibits affinity for ssDNA (Figure 4). Starting from 2.5 µM Gp204, a DNA smear could be seen on the agarose gel with increasing protein concentrations, indicating that Gp204 interacts with the ssDNA of phiX174. However, the observed shift was not as clear as the positive control, but this could be due to the lower protein concentration of Gp204 compared to the control where a concentration of 69 µM was used. Combining all results, it is hypothesized that Gp204 functions as a transcriptional regulator, thereby changing the transcriptome of ESBL098.

**Figure 4:**
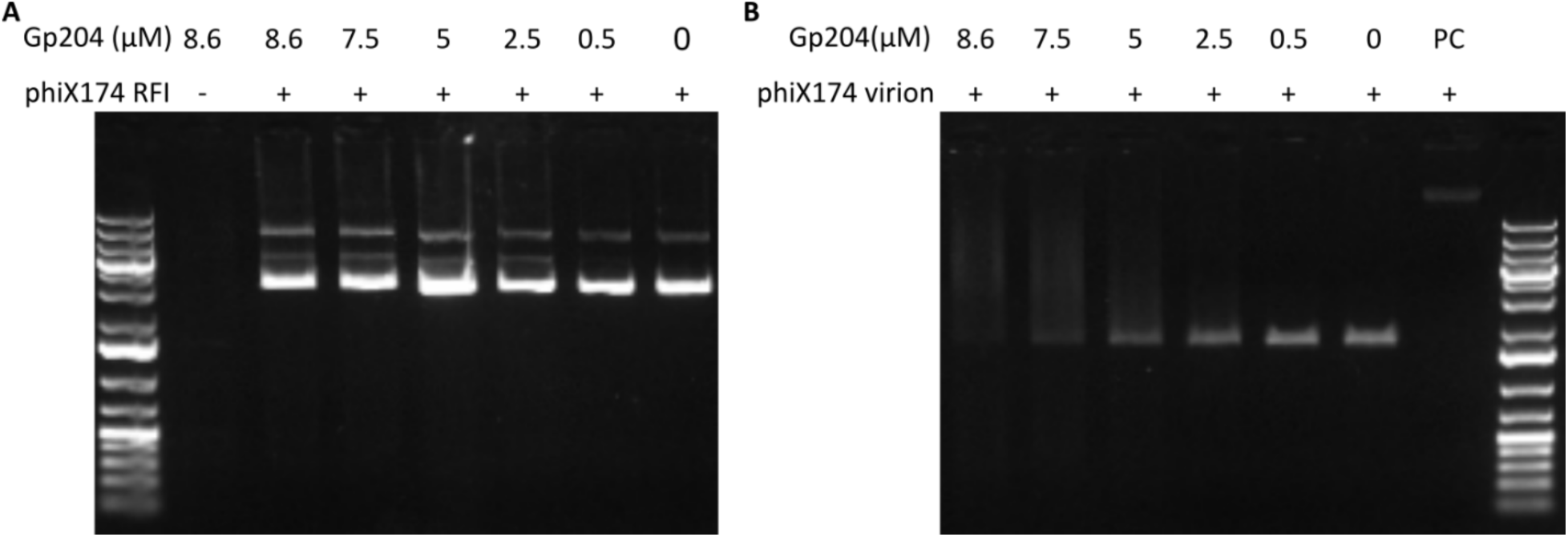
Gp204 binds ssDNA. EMSA of decreasing concentrations of Gp204 in combination with A) dsDNA of phiX174 (5.7 nM) or B) ssDNA of phiX174 (5.7 nM). The ssDNA binding protein Gp32 (69 µM) encoded by phage T4 is used as the positive control (PC).

Gp204 alters the transcriptome of *E. coli* ESBL068 and is renamed Htm, Host transcription modulator To evaluate the impact of Gp204 on the ESBL098 transcriptome, a differential gene expression analysis was performed comparing gene expression levels between ESBL098 transformed with the empty vector pPS26 and pPS26::*gp204* after being induced with rhamnose for 30 minutes. A principal component analysis revealed that the two conditions could be distinguished from each other, although there was more variation in the control samples. In total, 738 genes were identified with an absolute log2 fold change above 2 and an adjusted p-value smaller than 0.05, indicating a significant impact of Gp204 expression on the transcriptome. Of these, 586 genes were upregulated, and 152 were downregulated (Figure 5A). To determine the biological processes involved, KEGG pathway enrichment analysis was performed for both the significantly up- and downregulated genes. The analysis revealed that the downregulated genes were primarily involved in flagellar assembly and two-component systems (Figure 5B). The flagellar assembly pathway included structural genes for the MS/C ring, the rod, P/L ring, and the hook, suggesting that flagella formation is likely reduced. The downregulated two-component systems affected various processes, such as citrate metabolism and capsule formation. The citrate/succinate antiporter *citT* and citrate lyase genes *citCDEFXG* were downregulated, though *citC* and *citG* were not statistically significant. Additionally, the last three genes of the operon encoding the group 4 capsule polysaccharide - *gfcE, etk,* and *etp* - were also significantly downregulated.

**Figure 5:**
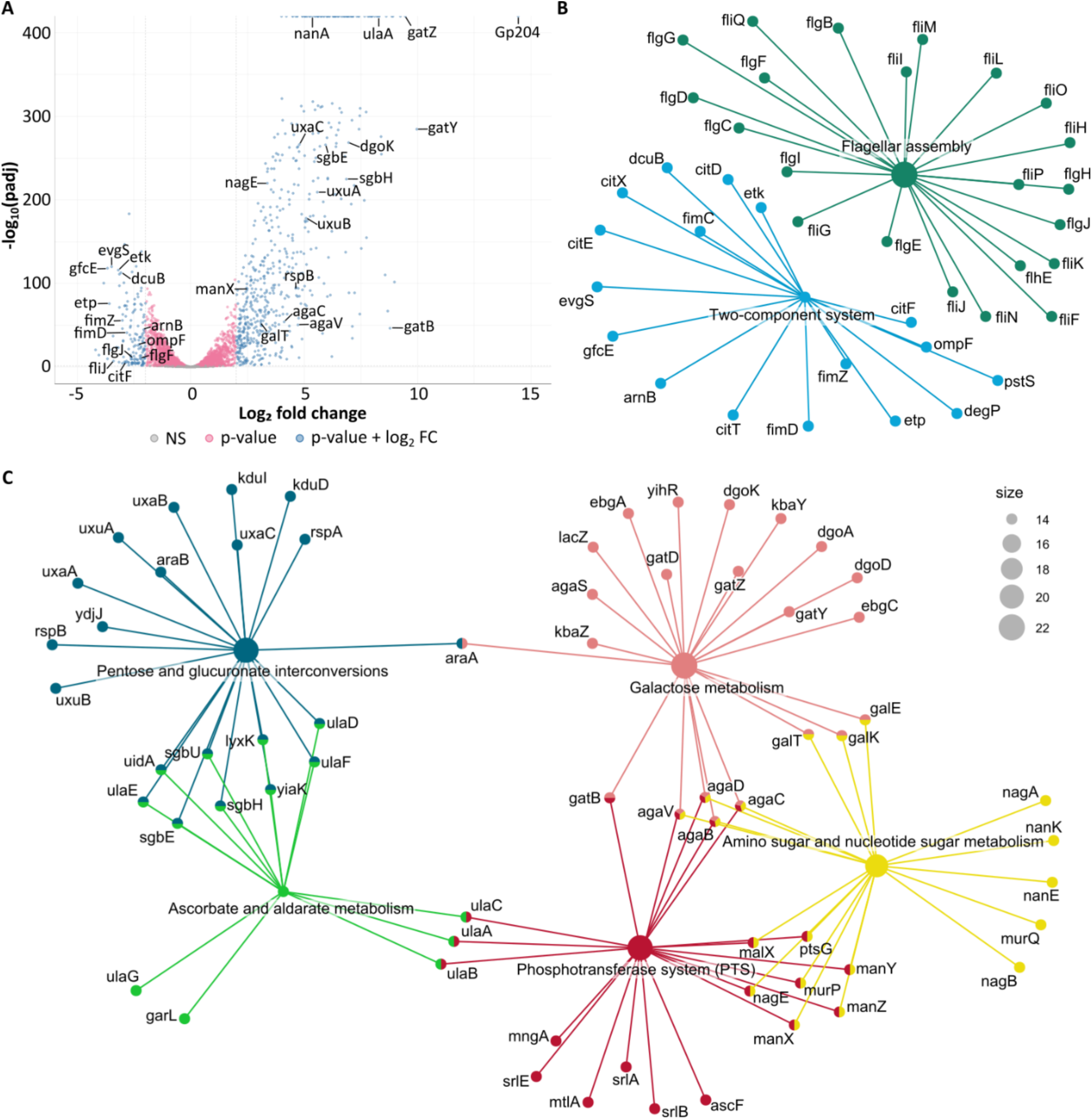
Gp204 causes differential expression of genes involved in diverse pathways. A) Volcano plot showing the up- and down-regulation of genes upon expression of Gp204. B-C) Interaction plot of differentially expressed genes involved in the most significantly B) down- and C) up-regulated pathways.

The five most significantly enriched pathways in the upregulated genes are galactose metabolism, pentose and glucuronate interconversions, ascorbate and aldarate metabolism, phosphotransferase systems (PTS), and amino sugar and nucleotide sugar metabolism (Figure 5C). Upregulated genes in the first two pathways enhance the production of various precursors, such as D-glyceraldehyde-3-phosphate and D-ribulose-5-phosphate, which are further metabolized in the pentose phosphate pathway. Genes involved in ascorbate and aldarate metabolism promote the formation of D-xylulose-5-phosphate, which enters the pentose and glucuronate interconversion pathway and eventually the pentose phosphate pathway. This pathway is essential for the formation of D-ribose-5-phosphate, which in turn is a precursor for the metabolism of purines, pyrimidines, and histidine. Upregulation of PTS results in increased phosphorylation of glucose- and mannose-based sugars, while the genes in amino sugar and nucleotide sugar metabolism lead to an increase in GlcNAc-6-phosphate and ManNAc-6-phosphate, which are involved in nucleotide sugar biosynthesis. Less significantly enriched pathways included phenylalanine metabolism, benzoate degradation, fatty acid degradation, butanoate metabolism, tryptophan metabolism, and the degradation of valine, leucine, and isoleucine. All these pathways increase acetyl-CoA, which can enter the citrate cycle. Through the citric acid cycle, energy supply can be increased. Lastly, there is an increase in L-tryptophan production, and the complete quorum-sensing pathway, except for *luxS*, is upregulated. Taken together, these Gp204-induced transcriptomic changes could benefit AV110 during phage infection by facilitating amino acid and nucleotide synthesis and increasing the available energy required to produce new phage progeny. These results also suggest that Gp204 potentially functions as a transcriptional regulator with a broad effect on the host transcriptome, therefor the protein was renamed Host transcription modulator (Htm).

## Discussion

When phages reprogram their host during infection, this process also involves reprogramming the host transcriptome. In this study, we characterized the activity of the *Mosigvirus* AV110-encoded protein Gp204 (Htm) as a potential transcriptional regulator and its effects on the transcriptome of ESBL *E. coli.* We demonstrated that recombinant expression of Htm inhibits the growth of all tested ESBL/AmpC *E. coli* strains. This inhibition is characterized by a strongly reduced growth rate, DNA condensation, and the differential expression of 738 genes, corresponding to 15% of the ESBL098 coding sequences. However, no increase in cell death was observed. The combination of a reduced growth rate with the absence of cell death could indicate that the effect is bacteriostatic rather than bactericidal, and cells could grow again upon removal of Htm.

Htm harbors the conserved AAA+ ATPase domain. This domain is present in proteins encoded by bacteria, eukaryotes, archaea, and viruses, and serves various functions (22). During phage infection, proteins carrying AAA+ ATPase domains are involved in DNA replication, DNA packaging into the capsid, and are part of the BREX phage defense system (28–30). To perform the different functions, modifications are made to the conserved domain structure (22). For example, in the transcriptional activator PspF, two loops are inserted into the ATPase domain with a conserved motif to facilitate the interaction with σ^54^ and activate transcription of the *psp* operon in *E. coli* (31). While the insertion detected in the conserved domain of Htm does not contain the same motif as PspF, it is likely involved in a protein-specific interaction that is important for its function. Additionally, a mutation was observed in the Walker B motif of the AAA+ ATPase domain, where the conserved aspartic acid and glutamic acid residues were replaced with two aspartic acid residues. A similar substitution in PspF resulted in increased ATP binding but a significant decrease in ATPase activity to 4% of the wild-type activity (32), suggesting that Htm’s ATPase activity might be similarly reduced. In related bacterial enhancer-binding proteins (bEBPs), which regulate transcription by activating the σ^54^-bound RNA polymerase, ATPase activity repositions two domains of σ^54^, thereby promoting the formation of the open complex and activating transcription (33). Thus, we propose that Htm could have a similar function.

In contrast to most bEBPs, Htm lacks the C-terminal DNA-binding domain that binds to enhancers upstream of the promoter, explaining its lack of affinity for dsDNA. However, it could bind ssDNA *in vitro*. While most transcriptional regulators bind to dsDNA, they can also interact with ssDNA, as was shown for the phage LUZ7 protein Drc (34). A model was proposed in which Drc binds to promoters on the LUZ7 genome that are locally unwound, recruiting the RNA polymerase and initiating transcription of middle genes (34). Htm could potentially bind to the -12 and -11 bases of the promoter sequence, which become unwound after the σ^54^-RNA polymerase complex binds in the closed conformation (35). This, together with the interaction of Htm with the sigma factor or RNA polymerase, could be sufficient to allow Htm to bind initially. Due to the use of the phiX174 genome as the DNA source, the binding between Htm and the ssDNA is most likely non-specific. To investigate specific binding, the ESBL098 and AV110 genomes could be used in ChIP-seq, which would simultaneously allow the determination of specific binding sites, if any, and indicate potential modulating activity of Htm on phage transcription. Demonstrating the interaction between Htm and the RNA polymerase by, for example, a mobility shift assay or a pull-down assay would provide further evidence for Htm’s potential role as a transcriptional regulator.

The N-terminal domain of Htm showed structural similarity to the C-terminal of NusG. This domain is known to bind both the transcription terminator ρ and the ribosomal protein S10. By binding ρ, NusG enhances its activity and thereby promotes transcription termination (26). The binding of the ribosomal protein S10 directly links transcription and translation (27). The recruitment of ribosomes to the site of active transcription could benefit the phage by enhancing and accelerating the production of host proteins required for phage infection. However, the interaction between Htm and ρ and/or rS10 still must be confirmed. In phage λ, NusG was shown to enhance transcription antitermination during lytic infection through the interaction with λN, NusA and rS10, thereby preventing binding of ρ (36, 37). Based on the experimental results and the structural similarity of Htm to bacterial enhancer-binding proteins and the C-terminal domain of NusG, we hypothesize that Htm interacts with the σ^54^-RNA polymerase complex and binds locally unwound DNA to activate transcription. Through its N-terminal domain, it may competitively bind to the ribosomal protein S10 or ρ to link transcription to translation or to terminate transcription, respectively.

The bacterial pathways affected by Htm expression suggest a decreased cell motility and increased levels of precursors of the pentose phosphate pathway for the metabolism of purines, pyrimidines, and histidine, as well as increased nucleotide sugar biosynthesis and energy supply through an increase in acetyl-CoA. It is well known that phage infection has a strong effect on the host transcriptome, the proteome, and metabolome (38). Multiple studies have investigated pathways that are either up- or downregulated upon phage infection, showing that the effect can be highly phage-dependent. In a study examining the metabolic changes induced by infection of six *Pseudomonas* phages, an increase in nucleotide sugar metabolism was the only conserved metabolic change (16). It was also shown that the effect of phage infection on the host transcriptome depends on growth conditions (13). Most studies focus on the effect of phage infection on the host, with only limited studies on the impact of individual phage proteins on the host transcriptome, proteome, and metabolome. In one study, the authors investigated the effect of λ Orf73, a TraR homolog, and found that it reduces cell growth and inhibits the synthesis of translation machinery, while increasing phage yield during lytic infection (18). Another example is AppY, encoded by the *E. coli* K-12 prophage DLP12, which improved acid stress resistance and biofilm formation but also decreased cell motility (39). Similar to AppY, Htm downregulated the expression of genes in the *fli* and *flg* operons, which encode the structural components of the flagella (39). As a result, cell motility is expected to decrease. This can be beneficial during phage infection by redirecting resources from the energetically costly flagellum formation and flagellum-driven motility to increased progeny production. The combination of up- and downregulated processes can contribute to an increased production of phage progeny by increasing the availability of amino acids, nucleotides, and energy. In conclusion, our results show that Htm likely functions as a transcriptional regulator to modulate the host transcriptome, thereby enhancing the availability of resources, such as nucleotides and energy, to produce phage progeny. These results provide new insights into the effect of an individual phage protein on host reprogramming during phage infection.

## Materials and Methods

### Bacteria, phages, and plasmids

The bacteria, phages, and plasmids used in this study are listed in **Error! Reference source not found.**. Bacteria were grown in LB, when necessary, supplemented with kanamycin (50 µg/ml) and incubated overnight at 37°C with shaking at 180 rpm.

### Cloning of Gp204

Gp204 was amplified from the AV110 genome with the addition of overhangs homologous to the vector and the vector was linearized with PCR using the primers listed in **Error! Reference source not found.** and the CloneAmp HiFi PCR Premix (Takara). The amplicon sizes were verified on a 1% agarose gel and the PCR products were purified using the GeneJet PCR Purification Kit (Thermo Scientific). The vector with the correct insert was constructed using the In-Fusion Snap Assembly Master Mix and transformed into Stellar^TM^ competent cells (Takara) via heat shock. After overnight incubation, the insert size was verified with PCR, and the plasmid was purified using the GeneJet Plasmid Miniprep Kit (Thermo Scientific). The purified plasmid was transformed into electrocompetent ESBL/AmpC *E. coli* strains or *E. coli* BL21 via electroporation.

### Growth inhibition assay

The growth-inhibitory effect of Gp204 in different ESBL/AmpC *E. coli* strains and *E. coli* Stellar^TM^ was assessed in biological triplicate. Tenfold dilution series were prepared from overnight cultures of the different strains transformed with pPS26::*gp204*. Each dilution was spotted on LA plates with kanamycin with or without 5 mM rhamnose. Plates were incubated overnight at 37°C.

### Growth curves

A growth curve of ESBL098 pPS26::*gp204* was constructed in biological triplicates by growing the culture to OD600 of 0.2. Next, the culture was twofold diluted in LB with kanamycin, with or without 5 mM rhamnose, to a final volume of 200 µl. Cultures were incubated in a 96-well plate at 37°C with OD600 measurements every 15 min for 24 h using the PowerWave XS microplate reader (BioTek).

### Time-lapse microscopy with and without live-dead staining

An overnight culture of ESBL098 pPS26::*gp204* was diluted 1/100, incubated for 1 h at 37°C, and then diluted to an OD600 of 0.01. The diluted culture was spotted onto a 1.2% agarose pad in LB with kanamycin and 5 mM rhamnose in a GeneFrame, then sealed tightly with a coverslip. The growth of three cells was followed for 25 h by acquiring phase-contrast images every 5 minutes using an inverted Olympus IX83 microscope with a 1.4 N.A. 100x oil-immersion objective and cellVivo incubator module set to 37°C. For live-dead microscopy, 1 µM propidium iodide was added to the agarose pad, and phase contrast and fluorescent images were acquired every 10 min.

### Fluorescence microscopy

An overnight culture of ESBL098 transformed with the empty vector or pPS26::*gp204* was diluted to an OD600 of 0.05 and grown for 30 min at 37°C. Next, the culture was grown for 1, 2, or 4 h in the presence of 5 mM rhamnose and 10 µg/ml Nile Red. Finally, cells were incubated with 100 µM OGDA and 5 µg/ml DAPI for 3 min at 37°C, after which the cells were harvested and fixed in 4.6% formaldehyde. After overnight incubation, cells were washed, resuspended in PBS, and spotted onto a 1.2% agarose pad in PBS. Fifteen images for each condition and time point were acquired using an inverted Olympus IX83 microscope with a 1.4 N.A. 100x oil-immersion objective.

### Protein localization

The location of Gp204 in the cell was determined by diluting an overnight culture of ESBL098 pPS26::*gp204* to an OD600 of 0.001 and incubating the diluted culture for 1 h at 37°C. Next, Gp204 expression was induced by the addition of 5 mM rhamnose and cells were grown for 4 h. Cells were stained with 100 µM OGDA, 1 µM Janelia Fluor® 646 HaloTag® Ligand, and 1 µg/ml DAPI, fixed in 4% PFA, and resuspended in PBS. Cells were spotted on a 1.2% agarose pad with ProLong Glass anti-fade mounting media and sealed with a coverslip. Images were acquired on an Abberior STED microscope (Abberior instruments) with a 60x 1.4 N.A. oil immersion objective. A matrix detector was used for the green 488 channel, a STED laser for the red 647 channel with 15 nm pixel size, and standard confocal imaging for the blue 405 channel.

### Protein expression and purification

A 5 ml overnight culture of BL21 pET28::*gp204* or BL21 pET28::*gp32* was added to 500 ml LB with kanamycin and grown at 37°C with shaking until OD600 of 0.8. Next, 0.5 mM IPTG was added and the culture was incubated overnight at 16°C with shaking. Cells were harvested by centrifugation at 4000 rpm for 30 min, resuspended in lysis buffer (20 mM Na2HPO4, 0.5 M NaCl, and 50 mM imidazole), and sonicated 8 times for 9 cycles at 80% power. Next, cell debris was removed through centrifugation at 10000 rpm for 20 min and the supernatant was filtered with a 0.45 µm filter and twice with a 0.22 µm filter. The protein was purified from the filtered supernatant using the His GraviTrap Purification Kit (Cytiva) and eluted in elution buffer (20 mM Na2HPO4, 0.5 M NaCl and 0.5 M imidazole). Using an Amicon Ultra-15 Centrifugal Filter Unit with a 10 kDa molecular weight cut-off (Millipore), the buffer was changed to TBS and the protein was concentrated. The presence of the correct band size was verified on a 4-12% NuPAGE^TM^ Bis-Tris SDS-PAGE gel (Invitrogen) and the protein concentration was determined with the Pierce Bradford Plus Protein Assay Kit (ThermoFisher Scientific).

### Electrophoretic mobility shift assay

A dilution series of the purified protein Gp204 was prepared in TBS. Next, 5 µl of each dilution was mixed with 2 µl 5x EMSA buffer (20 mM Tris pH 8, 1 mM MgCl2, 50 mM NaCl, 0.4 mM EDTA, 0.1 mM DTT, and 12.5% glycerol), 2 µl milliQ water, and 1 µl phiX174 RFI DNA (100 ng/µl; New England Biolabs) or phiX174 virion DNA (200 ng/µl; New England Biolabs). As a positive control, the T4 ssDNA binding protein Gp32 was added. As negative controls, mixtures were prepared without protein or DNA. The mixture was incubated at room temperature for 20 min, after which 2 µl 6x TriTrack DNA Loading Dye (Thermo Scientific) was added. The mixture was loaded on a 1% agarose gel with ethidium bromide and run for 30 min at 100 V. DNA bands were visualized using the Gel Doc XR+ (Bio-Rad).

### ESBL068 genome sequencing

The ESBL098 DNA was extracted using the Blood and Tissue Kit (Qiagen) according to the manufacturer’s instructions. Next, 150 ng DNA was prepared for sequencing using the Rapid Barcoding Kit 24 V14 (Oxford Nanopore Technologies). The prepared library was loaded on a MinION flow cell R10.4.1 (Oxford Nanopore Technologies) and sequenced for 29 h with the acquisition of fast5 files. The obtained reads were basecalled and demultiplexed using Guppy and filtered for a minimum quality of 12 and a minimum read length of 2000 bp with headcropping using Chopper (40). Quality control of the reads was performed with Nanoplot (40) and FastQC (41). Next, a hybrid genome assembly was performed by combining the obtained nanopore sequencing reads with Illumina sequencing data retrieved from ENA (Project PRJEB14641, Sample Accession SAMEA4058413, and Run Accession ERR1512441). An initial assembly was made using Trycycler (42) and further polished with Medaka (43) and Polypolish (44). Next, the genome was annotated with Prokka (45) with *Escherichia coli* specified as the genus and species.

### RNA sequencing

Three overnight cultures of ESBL098 transformed with either pPS26 or pPS26::*gp204* were diluted to an OD600 of 0.05 and incubated for 1 h at 37°C with shaking. Next, 5 mM rhamnose was added and the cultures were incubated for 30 min at 37°C with shaking before pelleting the culture by centrifugation at 8000 rpm for 5 min. The pellet was resuspended in RLT buffer (Qiagen) with 1% β-mercaptoethanol. Next, cells were lysed with ≤106 µm glass beads (Sigma-Aldrich) using the FastPrep-24 (MP Biomedicals), cooled on ice for 2 min and lysed again. The lysed cells were centrifuged for 2 min at 14000 rpm and the supernatant was mixed with an equal volume of 70% ethanol. The RNA was further purified using the RNeasy Mini Kit (Qiagen) with on-column DNase I digestion. The RNA concentration and RNA integrity number (RIN) were determined using Nanodrop and the RNA 6000 Nano Kit (Agilent) for the 2100 Bioanalyzer (Agilent), respectively. The RNA library was prepared using the NEBNext rRNA Depletion Kit (Bacteria) and the NEBNext Ultra II Directional RNA Library Prep Kit for Illumina (New England Biolabs) and sequenced on a NovaSeq 6000 platform (Illumina) by Amplexa Genetics.

### Di<erential gene expression analysis

RNA reads were trimmed and filtered to a minimum length of 60 bp with Trimmomatic. Next, an index of the ESBL098 pPS26::*gp204* genome was prepared and the reads were mapped against the index with hisat2 (46). The obtained SAM files were converted to BAM files with SAMtools (47). Read count files were obtained using featureCounts (48) requiring a minimum mapping quality of 10. Next, genes with fewer than ten reads in at least two samples were removed and a differential gene expression analysis with DESeq2 and a principal component analysis were performed in R v4.5.1. Genes with an adjusted p-value below 0.05 and an absolute log2 fold change higher than 2 were selected as significantly differentially expressed. To perform a KEGG pathway enrichment analysis, the gene IDs were converted to the corresponding *E. coli* K-12 MG1655 KEGG IDs (https://rest.kegg.jp/list/eco) and the analysis was performed for the up- and downregulated genes using the function enrichKEGG from the R package clusterProfiler v4.16.0. The relation between the different enriched pathways and differentially expressed genes was visualized using the function cnetplot from the R package enrichplot v1.28.4.

## Data availability

The raw RNA sequencing data and read counts per gene, the nanopore sequencing reads of ESBL098 and the assembled and annotated ESBL098 pPS26::*gp204* genome are available on https://data.mendeley.com/preview/85d9kpm7vr?a=9ab9d894-9f79-4c81-a06c-be5e5fe13394.

## Acknowledgements

This work was supported by the Independent Research Fund Denmark (2035-00112B).

## Author contributions

S.S.F. Van Overfelt: Conceptualization, Formal analysis, Investigation, Methodology, Visualization, Writing – original draft, Writing – review & editing

V. H. Mebus: Conceptualization, Investigation, Methodology, Writing – review & editing

J. Brewer: Conceptualization, Writing – review & editing

R. Lavigne: Conceptualization, Funding acquisition, Writing – review & editing

A. N. Sørensen: Conceptualization, Methodology, Supervision, Writing – review & editing

L. Brøndsted: Conceptualization, Funding acquisition, Methodology, Project administration, Supervision, Writing – review & editing

