## Supplementary information for "A phage-encoded host transcription modulator, Htm, reprograms the transcriptome of ESBL *Escherichia coli*"

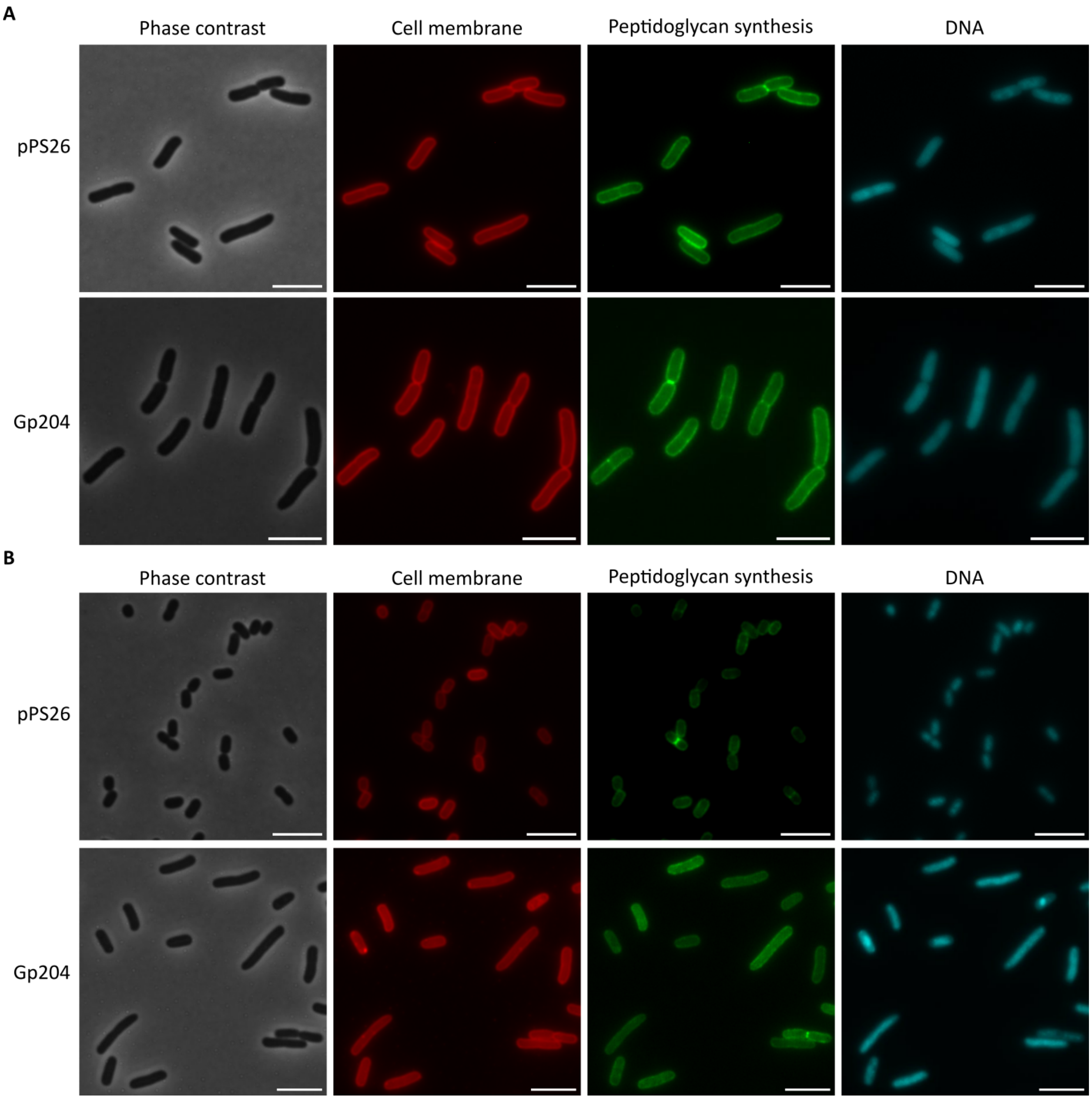


**Figure S1:** Phase contrast and fluorescence microscopy of ESBL098 transformed with the empty vector and pPS26::*gp204* after A) 1h and B) 4h of induction. The cell membrane, active sites of peptidoglycan synthesis, and the DNA are stained. Scale bar = 5 μm.

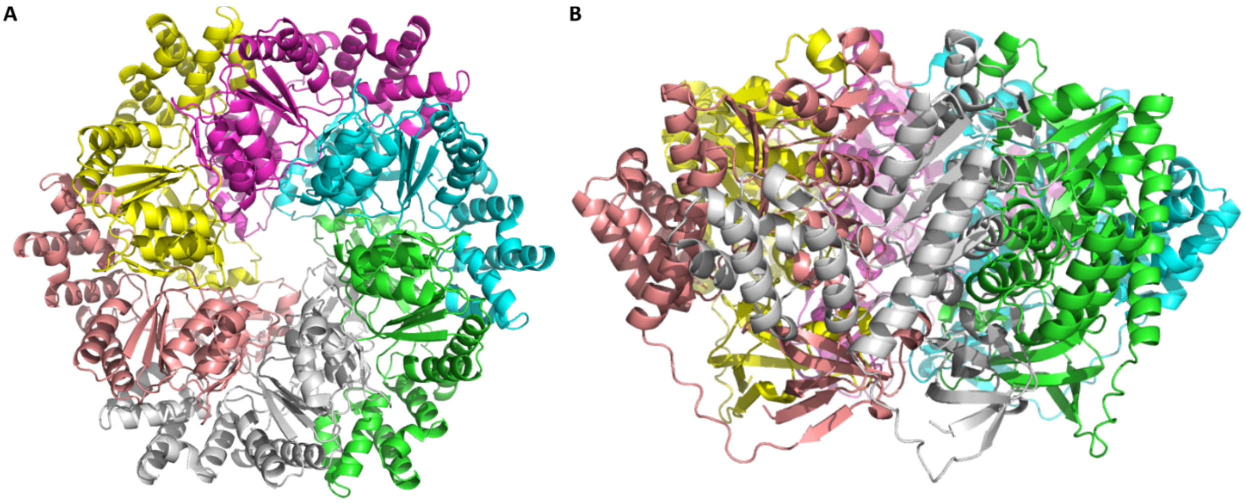


**Figure S2:** A) Top and B) side view of Gp204 assembled as a hexamer as predicted by AlphaFold 3.

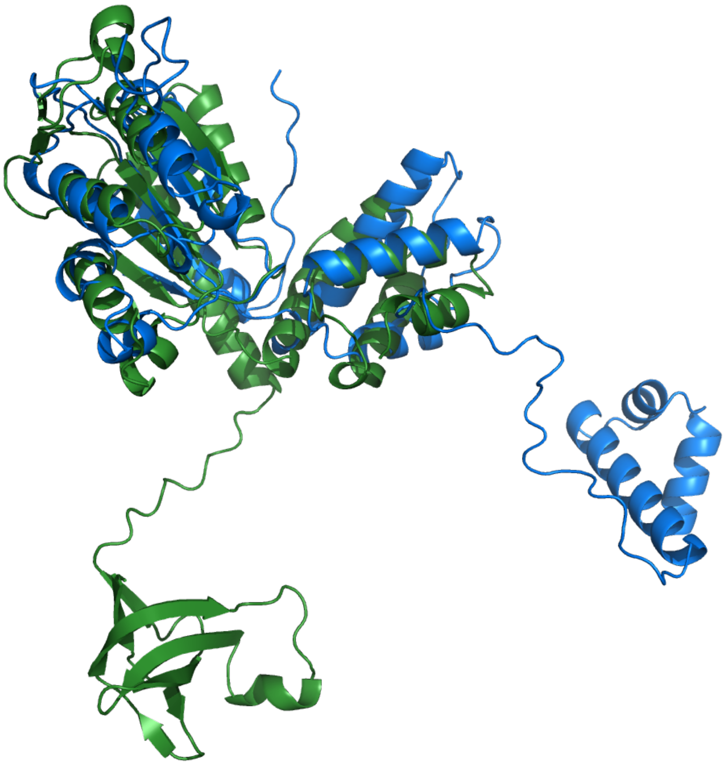

**Figure S3:** Overlay of the predicted structures of Gp204 (green) and PspF (blue, AF-P37344-F1-v6).

**Table S1:** Overview of the strains, phages, and plasmids used in this study.

| **Strains** | **Reference or source** |
|---|---|
| ESBL098 | (49) |
| ESBL032 | (49) |
| ESBL049 | (49) |
| ESBL054 | (49) |
| ESBL080 | (49) |
| ESBL087 | (49) |
| ESBL115 | (49) |
| ESBL154 | (49) |
| *E. coli* Stellar$^{TM}$ cells | Takara Bio |
| *E. coli* BL21 | Thermo Scientific |
| **Phages** | |
| AV110 | (50) |
| **Plasmids** | |
| pPS26 | (51) |
| pPS26::*gp204* | (20) |
| pPS26::gp204-Halotag | This study |
| pET28::*gp204* | This study |
| pET28::*gp32* | This study |

**Table S2:** Overview of the primers used in this study for protein purification and protein localization.

| **Primer ID** | **Sequence (5' -> 3')** |
|---|---|
| AV110_204-pET28-Fw | GGATCCGAATTCGAGATGTTCAACGTTCAAATCAAAAAAG |
| AV110_204-pET28-Rv | AGCTTGTCGACGGAGTTATTTCAGTAAAGTTACTTCAGAG |
| T4-gp32-pET28_Fw | CCGCAAGCTTGTCGACGGAGATGTTTAAACGTAAATCTACTGC |
| T4-gp32-pET28_Rv | GTCGCGGATCCGAATTCGAGTTAAAGGTCATTCAAAAGGTC |
| pPS26_gp204-Halo_Fw | CACCTCAGCGTCGTGACTG |
| pPS26_gp204-Halo_Rv | TTTCAGTAAAGTTACTTCAGAGATTTCC |
| Halotag-gp204_Fw | GTAACTTTACTGAAAGGAGGAGGTGGAAGTCC |
| Halotag-gp204_Rv | CACGACGCTGAGGTGTCAGCCGGAAATCTCGAGC |

**Video S1:** Time-lapse microscopy of ESBL098 pPS26::*gp204* grown on LB supplemented with 5 mM rhamnose for 24 h. Scale bar = 10 μm.
